# Therapeutic ICOS blockade reduces T follicular helper cells and improves allergic airway disease

**DOI:** 10.1101/250217

**Authors:** F. I. Uwadiae, C.J. Pyle, S.A. Walker, C.M. Lloyd, J.A. Harker

## Abstract

Allergic asthma is a disease of chronic airway inflammation and remodelling, characterised by a dysregulated type 2 response and allergen-specific IgE. T follicular helper cells (T_FH_) are critical to antibody production and have recently been implicated in allergic airway disease (AAD) pathogenesis. The role of T_FH_ in established disease and the therapeutic potential of targeting them are however not fully understood. Using two aeroallergen driven murine models of chronic AAD, T_FH_ were first identified in the lung draining lymph nodes but with prolonged exposure were present in the lung itself. Sustained allergen exposure led to the accumulation of T_FH_, and concomitant development of germinal centre B cells. Blockade of Inducible T cell co-stimulator (ICOS) signalling during established AAD depleted T_FH_ without adversely affecting the differentiation of other CD4^+^ T cell subsets. This resulted in impaired germinal centre responses, reduced allergen specific IgE and ameliorated inflammation and airway hyper-responsiveness, including reduced pulmonary IL-13. T_FH_ did not however appear to produce IL-13 directly, suggesting they indirectly promote type-2 inflammation in the lungs. These data show that T_FH_ play a pivotal role in the regulation of AAD and that targeting the ICOS-L pathway could represent a novel therapeutic approach in this disease.

## INTRODUCTION

Allergic asthma is a disease of chronic airway inflammation and remodeling, associated with a dysregulated type 2 immune response. The disorder is driven by chronic exposure to aeroallergens, including house dust mite (HDM) and fungal spores, leading to production of type 2 cytokines, IL-4, IL-5 and IL-13, and the hallmark symptoms of allergen specific immunoglobulin E (IgE), eosinophilia and airway hyper-responsiveness (AHR)^1^. Traditionally, the disorder has been described as a T helper 2 (Th2) cell disease, as they produce type 2 cytokines which drive the pathophysiology. However, it is now clear that multiple other cells of the immune system can produce these cytokines and thus could play vital roles in the regulation of distinct asthma phenotypes^1^.

T_FH_ are a specialised subset of CD4^+^ T cells with a unique capacity to help B cells produce high affinity, isotype-switched antibodies and differentiate into memory B cells and plasma cells^2^. They are defined by expression of CXCR5, PD1, Bcl-6 and ICOS and reside within the B cell follicles of secondary lymphoid organs, including lymph nodes and the spleen^2-7^. T_FH_ differentiation is a multi-step process. Firstly, within the T cell zone, naive CD4^+^ T cells are presented antigen and receive co-stimulation via inducible T cell co-stimulator ligand (ICOS-L) from dendritic cells^8^. Next, pre-T_FH_, now expressing Bcl-6 and CXCR5, migrate towards the T/B cell zone border^9, 10^ where antigen presentation and co-stimulation is transferred to activated B cells^11, 12^. Fully differentiated T_FH_ are located in the B cell zone, within newly formed anatomical structures called germinal centres (GC)^10^. Here, GC B cells provide antigen and pro-survival ICOS-L stimulation to the T_FH_, which in return provide pro-survival and anti-apoptotic signals to the B cells^2, 9, 11^.

A handful of studies have now attempted to dissect the role of T_FH_ during AAD pathogenesis. T_FH_ have been identified in the lymph nodes and spleens of HDM sensitised and challenged animals^13-15^ and have been shown to be critical to the production of allergen specific IgE^16, 17^. T_FH_ generated in lung draining lymph nodes have been shown to become Th2 cells which migrate to the lungs on allergen challenge and promote exacerbated HDM mediated AAD^15^. In contrast, adoptive transfer of IL-21^+^ T_FH_ into non-sensitised animals failed to generate Th2 cells on HDM exposure, but drove airway eosinophillia^13^. Interestingly, during Th2 pathology T_FH_ have been shown to differentiate out of the Th2 lineage^18^ and even obtain effector functions related to other CD4^+^ T cell lineages^19^. While, Bcl-6 deficient CD4^+^ T cells, unable to differentiate into T_FH_, can more readily become lung resident Th2 cells and drive AAD^14^. Taken together, these studies imply T_FH_ are important to allergic disease but their precise role remains unclear, and the therapeutic potential of targeting them is untested.

In this study using two chronic allergen exposure models mimicking the repeated environmental exposures that allergic asthmatics experience, T_FH_ were readily identified within the secondary lymphoid organs and the lung tissue itself. ICOS is a co-stimulatory molecule required for T cell activation expressed on all CD4^+^ T cells following T cell receptor engagement^20^. T_FH_, but not other CD4 T cell subsets, require sustained ICOS/ICOS-L signalling after priming to maintain their phenotype^21, 22^. Fitting with this T_FH_ and GC B cells could be specifically reduced by ICOS-L blockade even after allergic disease had been established. Importantly therapeutic blockade of T_FH_ dampened hallmark features of allergic disease, including eosinophilia, AHR, allergen specific IgE production and reduced pulmonary IL-13. These findings suggest blocking ICOS/ICOS-L interactions and targeting T_FH_ responses can provide therapeutic benefit in established AAD.

## RESULTS

### Chronic allergen exposure generates local and systemic T_FH_ responses

Allergic asthma is a disease of chronic pulmonary inflammation driven by repeated low dose exposure to aeroallergens. To mimic the pathogenesis *in vivo* and determine if responses were antigen dependent, mice were exposed to two common aeroallergens; either house dust mite (HDM) or *Alternaria alternata* (ALT) 3 times a week for up to 5 weeks (Figure 1A).

**Figure 1:**
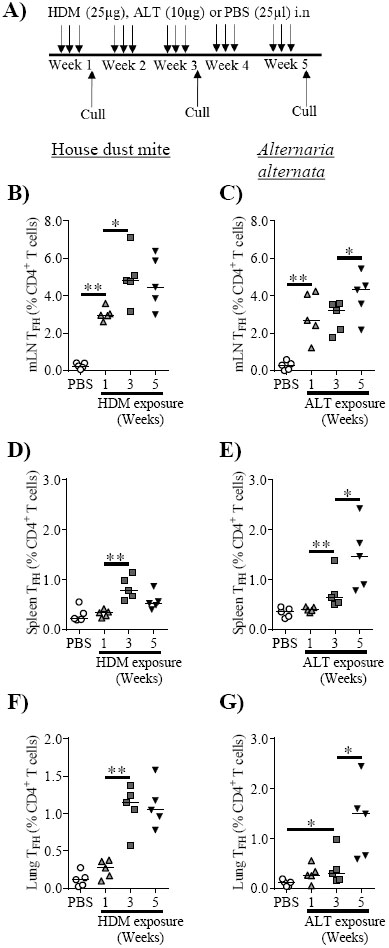
T _FH_ accumulate over time in the lungs and peripheral tissues. Adult female BALB/c mice were exposed to either 25 μg house dust mite (HDM), 10 μg *alternaria alternata* (ALT) or 25 μl phosphate buffered saline (PBS), 3 times a week for up to 5 weeks. Flow cytometry was used to determine the frequency of T_FH_ within cellular compartments following allergen exposure. T_FH_ were defined as CXCR5^+^PD1^+^Foxp3^-^CD4^+^ lymphocytes **A)** Experimental set up. Left panel – HDM, Right panel – ALT. **B-C)** mediastinal lymph nodes (mLN), **D-E)** spleen, **F-G)** lungs. Statistical significance was determined using a Mann Whitney U test. * *P<0.05*, \*\**p<0.01*, \*\*\**p<0.001*, n=5 per time-point. Representative data from 2 independent experiments.

T_FH_ (defined as CXCR5^+^PD1^+^Foxp3^-^CD4^+^) were observed in the lung draining mediastinal lymph nodes (mLN) of HDM or ALT treated animals after 1 week and very few were observed in mice only exposed to PBS (Figure 1B, C and Supplementary figure S1A). Continued allergen exposure further increased T_FH_ proportions in the mLN after 3 and 5 weeks. (Figure 1B, C and Supplementary figure S1A). Elevated T_FH_ frequencies were also found in the spleen, but only after 3 weeks of allergen exposure (Figure 1D, E and Supplementary figure S1B). Moreover, T_FH_ were observed at the site of inflammation, with HDM inhalation causing consistently elevated lung T_FH_ frequencies between weeks 3 and 5 (Figure 1F, Supplementary figure S1C). ALT lung T_FH_ were significantly increased at 3 weeks compared to PBS controls but only reached a frequency comparable to HDM after 5 weeks of exposure (Figure 1G, Supplementary figure S1C). Circulating T_FH_ were not observed in either model (Supplementary figure S2A-C). Taken together this showed that prolonged allergen exposure induced both tissue resident and systemic T_FH_ responses that increased in frequency over time. T_FH_ responses were established across all sites after 3 weeks of exposure.

### B cell responses and antibody development are preceded by T_FH_ responses during AAD

T_FH_ regulate antibody responses by directly interacting with activated B cells, driving the formation of germinal centres (GC), where isotype switching, affinity maturation and B cell maturation occurs^2^. GC B cells (defined as CD38^-^GL7^+^FAS^+^CD19^+^B220^+^) were absent in the mLN and lungs after 1 week of aero-allergen inhalation and were comparable to PBS treated mice (Figure 2A-D, Supplementary figure S3A). However, after 3 weeks of exposure to either aeroallergen, GC B cell frequencies were significantly elevated in the mLN, remaining consistently raised between weeks 3 and 5. (Figure 2A, C and Supplementary figure S3A). Lung GC B cells were also observed in HDM exposed animals at 3 weeks (Figure 2B and Supplementary figure S3B), while only after 5 weeks of ALT exposure were lung GC B cell frequencies significantly increased compared to PBS controls (Figure 2D and Supplementary figure S3B). Similarly, GC B cells were identified in the spleen of allergen exposed animals after 3 weeks (Supplementary figure S3C).

**Figure 2:**
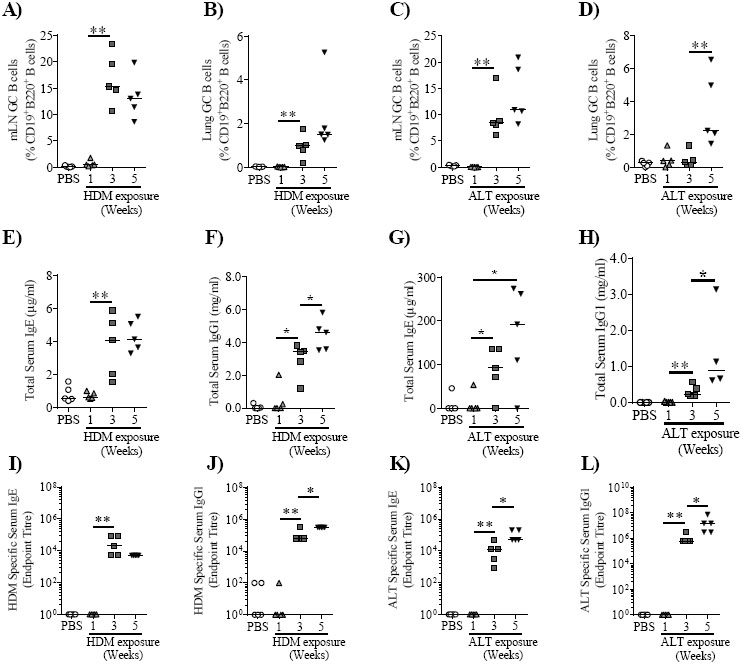
Allergen driven T _FH_ generation precedes germinal centre formation and antibody production. Adult female BALB/c mice were exposed to either 25 μg house dust mite (HDM), 10 μg *alternaria alternata (*ALT) or 25 μl phosphate buffered saline (PBS), 3 times a week for up to 5 weeks. Flow cytometry was used to determine the frequency of germinal centre (GC) B cells in the mLN and lungs. GC B cell were defined as CD38^-^GL7^+^FAS^+^CD19^+^B220^+^ lymphocytes and quantified. HDM study, **A)** mLN, **B)** Lungs, and ALT study, **C)** mLN, **D)** Lungs. Antibody within the serum was assessed by ELISA. **E)** HDM total IgE, **F)** HDM total IgG1, **G)** ALT total IgE, **H)** ALT total IgG1. Serum was titrated and allergen specific IgE and IgG1 was measured by ELISA. Endpoint titres are displayed. **I)** HDM-specific IgE, **J)** HDM-specific IgG1, **K)** ALT-specific IgE, **L)** ALT-specific IgG1. Statistical significance was determined using a Mann Whitney U test. * *P<0.05*, \*\**p<0.01*, \*\*\**p<0.001.* n=4-5 mice per time-point. Representative data from 2 independent experiments.

Allergen specific antibody, in particular IgE, is a hallmark feature of AAD. Total and allergen specific IgE and IgG1 were only detectable in allergen exposed mice after 3 weeks (Figure 2E-L). Sustained exposure to HDM did not further alter the concentration of total or specific IgE (Figure 2E, I), but did increase IgG1 production (Figure 2F, J). Total and ALT specific IgE and IgG1 were substantially increased at 5 weeks compared to 3 weeks (Figure 2G, H, K, L). Allergen specific antibodies were undetectable in the serum of PBS exposed animals (Figure 2I-L). This data shows the ability of chronic allergen exposure to generate local and systemic GC B cell responses and the subsequent emergence of allergen specific antibody titres that increase over time and are preceded by T_FH_ activity.

### ICOS/ICOS-L interactions are required to sustain T_FH_ during chronic AAD

T_FH_ require sustained signalling via ICOS to maintain their phenotype, and can be depleted by disrupting interactions between ICOS and its ligand, ICOS-L^9, 22^. To evaluate whether T_FH_ proportions could be reduced during established chronic allergic disease, mice were exposed to aeroallergens for 5 weeks and co-administered anti-ICOS-L antibody (α-ICOS-L) or an isotype control (2A3) in the last two weeks of allergen exposure (weeks 4 and 5). Mice were culled at the end of week 5 (Supplementary figure S4). 3 weeks of allergen exposure was sufficient to establish HDM and ALT driven allergic disease, including pulmonary inflammation, eosinophilia and AHR^23-25^. α-ICOS-L treatment substantially reduced mLN and lung T_FH_ populations after HDM exposure compared to 2A3 treated animals (Figure 3A-C). Similar results were observed with ALT (Figure 3A, D-E). Consistent with the reduced T_FH_ response, mLN and lung GC B cell responses induced by HDM inhalation were decreased in mice treated with α-ICOS-L compared to those given 2A3 (Figure 3F-H). In contrast, mLN GC B cells remained elevated in the ALT study (Figure 3F, I) and only lung proportions were reduced by α-ICOS-L treatment (Figure 3F, J). α-ICOS-L intervention caused decreased serum HDM-specific IgE (Figure 3K) while ALT specific IgE showed a trend to significant decrease (Figure 3L). Allergen specific IgG1 remained stable in both models after α-ICOS-L intervention (Figure 3M, N). Consistent with the reduced HDM specific IgE, but not ALT specific IgE, α-ICOS-L resulted in reduced HDM induced serum mast cell protease 1 (MCPT1) but had no impact on ALT induced MCPT1 (Supplementary figure S5). Taken together, this data shows that during chronic allergen exposure, T_FH_ can be successfully depleted using α-ICOS-L. The depletion altered GC B cell responses and allergen specific IgE but not IgG1.

**Figure 3:**
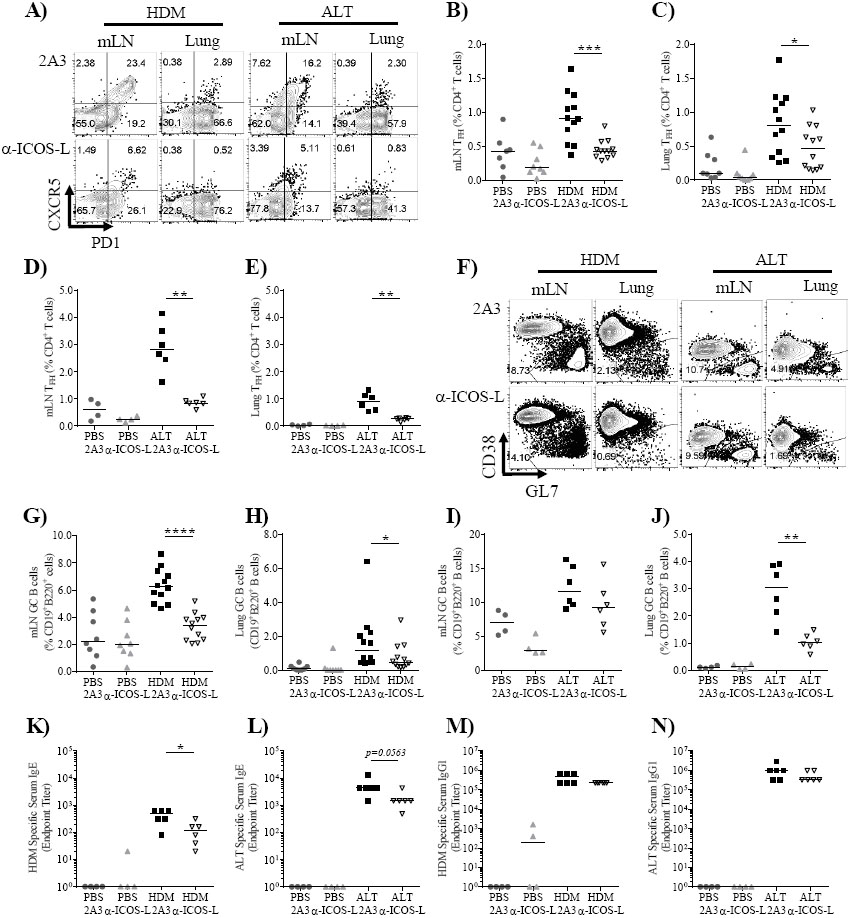
ICOS/ICOS-L interactions are required to sustain T _FH_ during chronic AAD. Adult female BALB/c mice were exposed (i.n) to 25 μg house dust mite (HDM), 10 μg *alternaria alternata (*ALT) or 25 μl phosphate buffered saline (PBS), 3 times a week for 5 weeks. From the start of week 4 mice were also administered 150 μg anti-ICOS-L (α-ICOSL) or isotype control (2A3) antibody (i.p) 3 times a week. Mice were culled at the end of week 5. **A)** Representative flow plots of mLN and lung T_FH_ following allergen and 2A3 or α-ICOS-L treatment, quantified in **B)** HDM mLN, **C)** HDM lung, **D)** ALT mLN, **E)** ALT lung. **F)** Representative flow plots of mLN and lung germinal centre (GC) B cells following allergen and 2A3 or α-ICOS-L treatment, quantified in **G)** HDM mLN, **H)** HDM lung, **I)** ALT mLN, **J)** ALT Lung. **K-N)** Serum was titrated and allergen specific antibody measured by ELISA. Endpoint titres are displayed. **K)** HDM specific IgE, **L)** ALT specific IgE, **M)** HDM specific IgG1, **N)** ALT specific IgE. Statistical significance was determined using a Mann Whitney U test. * *P<0.05*, \*\**p<0.01*, \*\*\**p<0.001.* HDM data shown is pooled from two independent experiments, n=8-12. ALT data is based on one study, n=4-6.

### α-ICOS-L blockade reduces pulmonary inflammation and airway hyperresponsiveness

Airway hyperresponsiveness and inflammation are fundamental indicators of AAD progression; therefore, the impact of α-ICOS-L blockade on the global allergic disease phenotype was examined. Mice exposed to HDM or ALT in combination with 2A3 displayed elevated cell numbers in the lung compared to PBS treated mice (Figure 4A, B). α-ICOS-L administration reduced aeroallergen induced cellular infiltration into the lungs (Figure 4A, B). Total lung eosinophils were reduced with allergen and αresulted in reduced airway resistance-ICOS-L co-administration (Figure 4C, D), however, the proportions of lungs eosinophils were unchanged (Supplementary figure S6).

**Figure 4:**
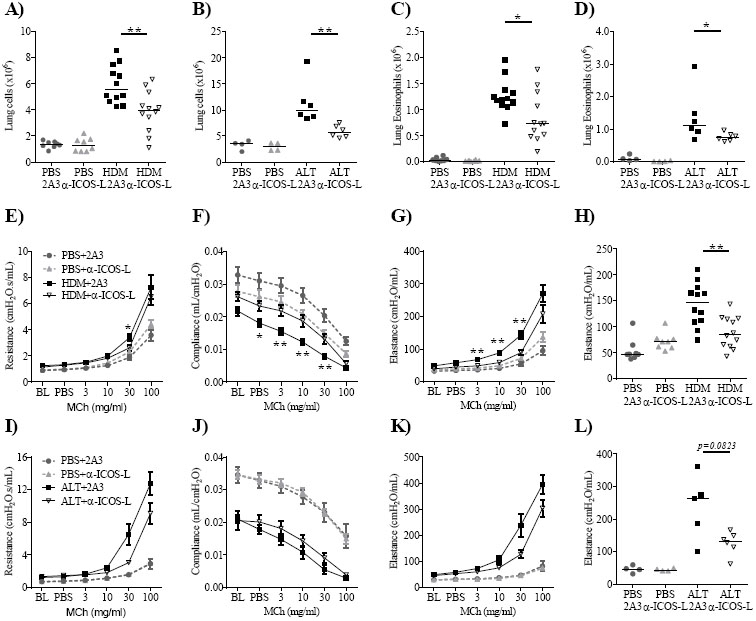
Therapeutic ICOS-L blockade improves chronic allergic airway disease. Adult female BALB/c mice were exposed (i.n) to 25 μg house dust mite (HDM), 10 μg *alternaria alternata (*ALT) or 25 μl phosphate buffered saline (PBS), 3 times a week for 5 weeks. From the start of week 4 mice were also administered 150 μg anti-ICOS-L (α-ICOSL) or isotype control (2A3) antibody (i.p) 3 times a week. Mice were culled at the end of week 5. Total number of cells in the lung, **A)** HDM study, **B)** ALT study. Total number of lung eosinophils, **C)** HDM study, **D)** ALT study. **E-H)** Airway hyperresponsiveness was measured by exposing mice to 0-100mg/ml methacholine (MCh) using the flexiVent system during HDM induced allergic airway disease. **E)** airway resistance, **F)** airway compliance and **G)** airway elastance, **H)** Quantification of airway elastance at 30mg/ml MCh. **I-L)** Airway hyperresponsiveness was measured during ALT induced allergic airway disease. **I)** airway resistance, **J)** airway compliance and K) airway elastance, **L)** Quantification of airway elastance at 30mg/ml MCh. Curves display mean±SEM. * = statistical difference between allergen+2A3 and allergen+α-ICOS-L. Statistical significance was determined using a Mann Whitney U test. * *P<0.05*, \*\**p<0.01*, \*\*\**p<0.001.* HDM data shown is pooled from two independent experiments, n=8-12. ALT data is based on one study, n=4-6.

AHR was evaluated by exposing mice to increasing doses of methacholine (MCh). Allergen exposure induced AHR, characterised by raised airway resistance and elastance, and reduced compliance compared to PBS controls (Figure 4E-L). Co-administration of HDM and α-ICOS-L resulted in reduced airway resistance and elastance and increased compliance, indicative of improved lung function compared to HDM and 2A3 treated animals (Figure 4EG). While α-ICOS-L treatment substantially decreased ALT induced airway resistance and elastance (Figure 4I, K) there was no impact on airway compliance (Figure 4J). In both models, large changes in airway elastance were observed (Figure 4H, L). Despite this, aeroallergen and α-ICOS-L administration had no impact on goblet cell hyperplasia (Supplementary figure S7A-C), collagen deposition (Supplementary figure S7D-F) or airway smooth muscle hyperplasia and hypertrophy (Supplementary figure S7G-I). However, taken together the data shows that therapeutic administration of α-ICOS-L after disease establishment improved airway inflammation and lung function.

### α-ICOS-L improves disease by reducing cellular inflammation rather than targeting Th2 cells or ILC2s

To assess the mechanism by which α-ICOS-L treatment could be facilitating effects on pulmonary inflammation and AHR, cytokine and chemokine secretion into the lungs was analysed. HDM or ALT treated animals administered 2A3 had increased IL-13 in their lungs compared to PBS exposed animals (Figure 5A, B). Therapeutic α-ICOS-L in combination with HDM reduced IL-13 in the lungs compared to 2A3 treated mice (Figure 5A) but this change was not seen during ALT exposure (Figure 5B). Similar trends were observed with lung IL-17A (Figure 5C, D). Allergen treatment induced IL-5 (Figure 5E, F) and eotaxin-2 (Figure 5G, H), but neither were altered by α-ICOS-L administration (Figure 5E-H). Thus, the HDM model appeared to be more susceptible to modulation by therapeutic α-ICOS-L treatment than the ALT model.

**Figure 5:**
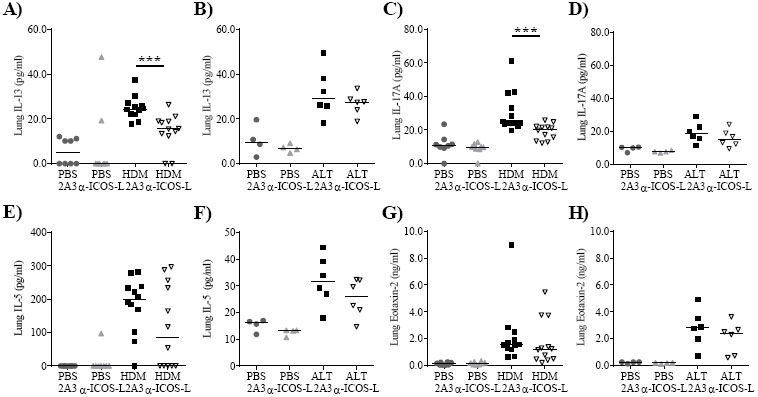
Therapeutic ICOS-L blockade reduces house dust mite induced pulmonary IL-13 and IL-17A. Adult female BALB/c mice were exposed (i.n) to 25 μg house dust mite (HDM), 10 μg *alternaria alternata (*ALT) or 25 μl phosphate buffered saline (PBS), 3 times a week for 5 weeks. From the start of week 4 mice were also administered 150 μg anti-ICOS-L (α-ICOSL) or isotype control (2A3) antibody (i.p) 3 times a week. Mice were culled at the end of week 5. Pulmonary cytokines and chemokines within the lung were measured by ELISA. IL-13 **A)** HDM study, **B)** ALT study. IL-17A **C)** HDM study, **D)** ALT study. IL-5, **E)** HDM Study, **F)** ALT study. Eotaxin-2, **G)** HDM study, **H)** ALT study. Statistical significance was determined using a Mann Whitney U test. * *P<0.05*, \*\**p<0.01*, \*\*\**p<0.001.* HDM data shown is pooled from two independent experiments, n=8-12. ALT data is based on one study, n=4-6.

Given the alteration in pulmonary IL-13 secretion and the importance of IL-13 to allergic disease pathogenesis, Th2 cells were studied in both models. HDM exposure increased the frequency of IL-13^+^ CD4^+^ T cells and this was unchanged by α-ICOS-L intervention (Figure 6A, B). However, consistent with decreased pulmonary inflammation (Figure 4A), total numbers of IL-13^+^ CD4^+^ T cells showed a trend towards reduction with α-ICOS-L treatment (Figure 6C). However, in the ALT study IL-13^+^ CD4^+^ T cells were reduced by α-ICOS-L treatment both by proportion and total number (Figure 6A, D-E). Similar trends were observed for IL-17A^+^ CD4^+^ T cells which were strongly induced upon HDM exposure and only decreased in total number consistent with the fall in overall pulmonary inflammation (Supplementary figure S8A-C). IL-17A^+^ CD4^+^ T cells were not significantly induced by ALT inhalation and were unchanged by α-ICOS-L intervention (Supplementary figure S8A, D-E).

**Figure 6:**
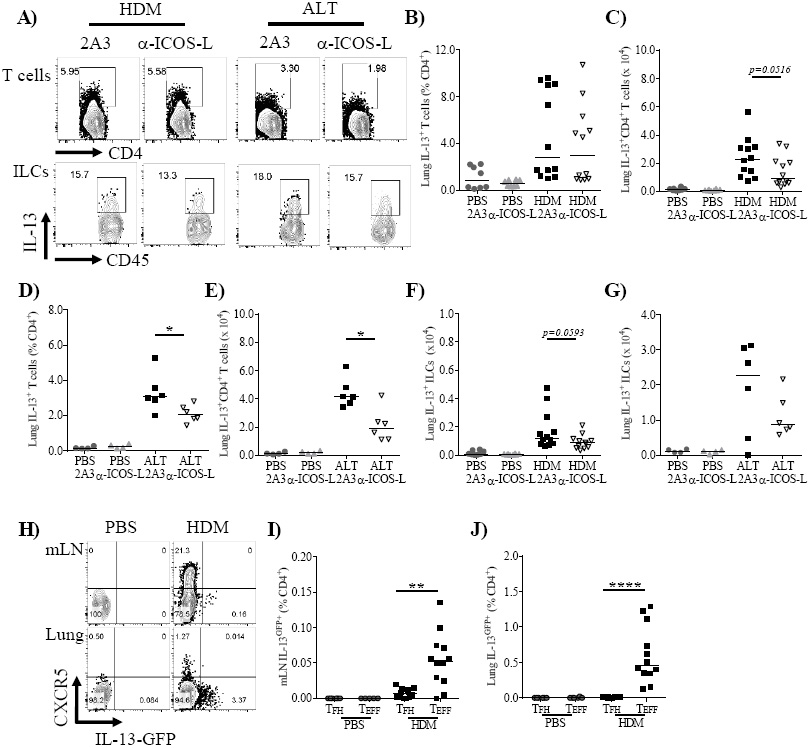
IL-13^+^ CD4^+^ T cells and ILCs are not directly targeted by ICOS-L blockade. **A-G)** Adult female BALB/c mice were exposed to either 25 μg house dust mite (HDM), 10 μg *alternaria alternata (*ALT) or 25 μl phosphate buffered saline (PBS), 3 times a week for up to 5 weeks. Flow cytometry was used to determine the frequency of lung cellular populations. **A)** Representative gating of IL-13^+^ CD4^+^ T cells and ILCs following allergen and 2A3 or α-ICOS-L treatment. **B)** Frequenc**y** IL-13^+^ CD4^+^ T cells – HDM study, **C)** Total IL-13^+^ CD4^+^ T cells – HDM study, **D)** Frequency of IL-13^+^ CD4^+^ T cells – ALT study, **E)** Total IL-13^+^ CD4^+^ T cells – ALT study, **F)** Total IL-13^+^ ILCs – HDM study, **G)** Total IL-13^+^ ILCs – ALT study. HDM data shown is pooled from two independent experiments, n=8-12. ALT data is based on one study, n=4-6. **H-J)** Adult female IL-13^GFP^ reporter mice were exposed (i.n) to 25 μg HDM or 25 μl PBS, 3 times a week for 3 weeks. Flow cytometry was used to determine the frequency of IL-13GFP,^+^ cells. **H)** Representative flow plots of CXCR5^+^ T cells and IL-13-GFP^+^ cells in PBS and HDM treated animals, pre-gated on CD4^+^CD3^+^CD44^hi^CD62L^-^ lymphocytes. Quantification of IL-13^GFP+^ T_FH_ (CXCR5^+^PD1^+^CD4^+^CD44^hi^CD62L^-^) and T_EFF_ cells (CXCR5^-^CD4^+^CD44^hi^CD62L^-^). I) mLN, J) Lung tissue. Data shown is pooled from two independent experiments, n=8-12. Statistical significance was determined using a Mann Whitney U test. * *P<0.05*, \*\**p<0.01*, \*\*\**p<0.001.*

ILC2s are a major producer of IL-13 during aeroallergen driven allergic disease^26^. HDM or ALT treatment along with 2A3 resulted in increased numbers of IL-13^+^ ILCs compared to PBS treated controls, but ILC2s were not affected by ICOS-L blockade (Figure 6A, F-G, Supplementary figure S9A, B). IL-17A^+^ ILCs were also analysed and were found unchanged by ICOS-L blockade during either ALT or HDM exposure by proportion or total number (Supplementary figure S9C-G). Collectively, these data suggest that α-ICOS-L antibody can therapeutically relieve established AAD by reducing pulmonary inflammation and AHR but not by directly targeting Th2 cells or ILCs.

T_FH_ have been shown to accumulate and become dysregulated during sustained antigen exposure^27, 28^. Given that T_FH_ were reduced together with secreted IL-13 and IL-13^+^ CD4^+^ T cells, whether T_FH_ were capable of producing IL-13 was examined. Using IL-13^GFP^ reporter mice, CXCR5 expression was found to be separated from IL-13^GFP^ expression in CD4^+^ T cells within the mLN and lungs (Figure 6H). Consistent with this the IL-13^GFP+^ cells were identified within the non-T_FH_ T effector (T_EFF_) population but not in the T_FH_ population with the mLN and lungs (Figure 6I, J). Thus, the reduction of T_FH_ was not directly responsible for the decreased IL-13 observed during HDM driven allergic disease. Despite this, the loss of T_FH_ during chronic AAD was critical for the impaired humoral response and may be indirectly responsible for the improved AAD. Overall the data presented here indicates α-ICOS-L treatment to be a beneficial intervention targeting humoral immunity and other hallmark symptoms of AAD (summarized in Supplementary Figure S10).

## DISCUSSION

AAD is commonly characterised by chronic exposure to aeroallergens, resulting in Th2 biased lung inflammation and dysregulated humoral immunity^1^. Here we sought to examine the importance of T_FH_ in the pathology of AAD and antibody mediated immunity using two clinically relevant aero-allergens. As expected, chronic allergen exposure resulted in the progressive development of AAD, including the production of IgE. T_FH_ developed over time, both systemically and locally, and were associated with the presence of GC B cells. Therapeutic administration of ICOS-L blocking antibodies interrupted T_FH_ responses, decreased humoral immunity and improved hallmark features of AAD.

In this study, we showed that T_FH_ populations were generated in the peripheral lymph nodes and with prolonged allergen challenge could be detected locally within the lungs. In previous acute studies T_FH_ have been shown to peak between 7 and 14 days post infection or protein vaccination, declining as antigen availability decreases^29, 30^. Here T_FH_ accumulated over time with repeated allergen exposure, consistent with other chronic disease models, such as repeated protein immunisations^31^, HIV^32^ and chronic LCMV infection^27, 28^. This observation fits with findings that sustained antigen and thus continuous TCR stimulation favours T_FH_ differentiation and results in more T_FH_ in chronic settings than acute^27, 28, 31^.

Chronic allergen exposure established a local lung T_FH_ population. Local T_FH_ have been observed during murine allergic disease^13^ and in other chronic human diseases, including within nasal polyps during chronic rhinosinusitis^33^, the synovial tissue during rheumatoid arthritis^34^ and Hepatitis B^35^ and C infected liver tissue^36^. However, we show for the first time that lung T_FH_ are found alongside lung GC B cells. Importantly isolated clusters of lymphoid cells are also present in the asthmatic lung and have been shown to be larger than in healthy controls^37^, implicating a role in pathology. This indicates that T_FH_ are likely present and dysregulated within the lungs of asthmatic patients and they may contribute to disease severity, through both regulating antibody production and Th2 associated pathology.

T_FH_ require sustained ICOS/ICOS-L signalling via phosphoinositol 3 kinase (PI3K) after priming to maintain their phenotype^22^. Common variable immunodeficiency patients with genetic defects in the ICOS gene fail to generate T_FH_^38^, while T_FH_ accumulate in *Roquin* mutants which overexpress ICOS^39, 40^. As a result blocking ICOS/ICOS-L interactions after T cell activation has been extensively used as a tool in multiple acute models to selectively deplete T_FH_ when antigen is limiting^9, 22, 31^. Here we show that late therapeutic administration of α-ICOS-L during established allergic disease can also reduce T_FH_ even when antigen is readily available during ongoing chronic inflammation. Concomitant with reduced T_FH_, GC responses were impaired and this short intervention reduced HDM-specific IgE, although it did not alter IgG1 levels over the same timeframe. The half-life of IgE is short in comparison to other immunoglobulin isotypes^41^ and transferred Der p1 and Lol p 1 specific IgE declined rapidly over a 50-day period while IgG remained relatively stable^42^. Therefore, a longer α-ICOS-L administration protocol may be required to alter more potent fungal specific IgE or the more stable IgG1^24^. Nonetheless, consistent with altered HDM-specific IgE levels, serum MCPT1 was significantly reduced indicating the treatment resulted in reduced mast cell activation^43^. Critically T_FH_ are required for the generation of antibodies including allergen-specific IgE^16, 17^, therefore this combined with our observation of reduced HDM specific IgE after a short intervention, highlights the potential of this approach to abrogate IgE driven clinical symptoms of AAD.

Alongside its’ effects on humoral immunity α-ICOS-L intervention after disease establishment improved both HDM and ALT driven AHR. IL-13 is a potent inducer of AHR^44, 45^ and consistent with this HDM induced pulmonary IL-13 was reduced following α-ICOS-L treatment. According to the literature T_FH_ are the only CD4^+^ T cell subset affected by late administration of a-ICOS-L^9, 22^. However, depletion of T_FH_ was not directly responsible for this reduced IL-13 as T_FH_ do not appear to be a major IL-13 source, indicating that during chronic AAD, T_FH_ did not acquire a T_FH2_ phenotype.

ILC2s are the major innate producer of IL-13 during chronic AAD^26^, while Th2 cells are the major adaptive source^1^. Furthermore, human and mouse ILC2s have been shown to depend on ICOS/ICOS-L signalling for their homeostatic survival and ability to initiate AAD^46^. Despite this, in both allergen models studied here ICOS blockade did not significantly reduce IL-13^+^ ILC2s, indicating that during established AAD ILC2s do require ICOS/ICOS-L signalling to function. Likewise, HDM induced IL-13^+^ CD4^+^ T cells showed a trend to decrease only in numbers upon ICOS-L blockade, consistent with the overall fall in gross cellular inflammation. While, ALT induced IL-13^+^ CD4^+^ T cells were significantly reduced both by proportion and total number. Thus Th2 cells may represent an indirect target of a-ICOS-L. In combination with the observation that IL-13^+^ CD4^+^ T cells outnumber IL-13^+^ ILCs in established disease, this indicates that CD4^+^ T cells are the major source of IL-13 and are responsible for the reduction of IL-13 after ICOS-L targeting.

This study is the first to show that disrupting ICOS/ICOS-L interactions after the establishment of allergic disease is both capable of depleting germinal centre reactions and has the potential to be therapeutically beneficial. Previous work has focused on blocking ICOS or ICOS-L prophylactically, during the inception of allergic disease^47-49^ or during an exacerbation^47^ using less clinically relevant ovalbumin induced allergic disease models^47-49^ rather than with ongoing allergen exposure with clinically relevant aeroallergens. Delivery of aeroallergens such as HDM directly to the airways and lungs in a chronic fashion better replicates human disease compared to intraperitoneal, skin sensitisation or short sensitisation and challenge models used in T_FH_ studies to date^13, 15, 16^. Thereby, improving our understanding on the response and role of T_FH_ in AAD. As α-ICOS-L blockade during established disease targeted multiple facets of the disease it could be potentially advantageous compared to currently approved biological therapies, such as omalizumab (anti-IgE mAb)^50^, reslizumab (anti-IL-5 mAb)^51^ and mepolizumab (IL-5 antagonist)^52^, which generally favour one arm of the allergic response over another. Furthermore, blocking ICOS signalling has been proven to be safe and effective in two phase I clinical trials for systemic lupus erthymatoesus^53^.

The results of this study give new insight into the role of T_FH_ in chronic AAD. Importantly, we show that T_FH_ can be identified locally within the lungs in addition to secondary lymphoid organs. Critically, even during chronic allergen exposure ICOS-L is required for the maintenance of T_FH_ and implicates α-ICOS-L blockade as a useful therapeutic intervention. This could be particularly essential for patients with severe disease where steroid treatment alone is not sufficient to control symptoms.

## MATERIALS AND METHODS

### Mice

6-8 week old female BALB/c mice were purchased from Charles River Laboratories (UK) and IL-13^GFP^ reporter mice were gifted from the lab of Professor Andrew McKenzie, University of Cambridge. Mice were housed in IVCs and all procedures were approved by the Imperial College London Animal Welfare Ethical Review Body (AWERB) and the United Kingdom Home Office (Approval from both under project licence number 70/7463) and conducted in accordance with the Animals (Scientific Procedures) Act 1986. All animal experiments are compliance with the ARRIVE guidelines. Mice were anesthetised via inhalation of isoflurane and euthanized by intraperitoneal overdose of pentobarbitone or low dose pentoberbitone with ketamine, using exsanguination via a peripheral vein as a secondary means of confirmation.

### Induction of allergic airway disease and ICOS-L intervention

Mice were administered 25 μg house dust mite (HDM) extract *(Dermatophagoides pteronyssinus)* or 10 μg *Alternaria alternata* (ALT) reconstituted in 25 μl PBS intranasally (i.n.) 3 times a week for up to 5 weeks (Greer Laboratories, NC, USA; Citeq, Groningen, The Netherlands). Control mice were given 25 μl PBS. In blocking experiments, from week 4 onwards, mice were co-administered 150 μg anti-ICOS-Ligand (Clone: HK5.3, BioXCell, NH, USA) or isotype control (Clone: 2A3, BioXCell, NH, USA) antibody in 200 μl PBS via intraperitoneal (i.p.) injection 3 times a week for 2 weeks. Mice were culled at the end of week 5. All mice were harvested 18 hours after the final allergen dose.

### Tissue Processing

Lung cells were disaggregated by incubating chopped tissue at 37 ˚C for 45 minutes in complete media (RPMI with 10% fetal calf serum, 2 mM L-glutamine and 100U/ml Penicillin/Streptomycin) containing 0.15 mg/ml collagenase type D (Roche Diagnostics, Burgess Hill, UK) and 25 μg/ml DNase type 1 (Roche Diagnositics). Splenic, mediastinal lymph node (mLN) and lung cells were recovered by filtering through a 100 μM nylon sieve and washing in complete media. Lung and splenic cells were then treated with red blood cell lysis buffer (155 mM ammonium chloride, 10 mM potassium bicarbonate and 0.1 mM Disodium EDTA) for 5 mins, washed and resuspended in complete media. Bronchoalveolar lavage (BAL) was collected by washing the airways three times with 0.4ml PBS via a tracheal cannula. BAL cells were pelleted and resuspended in 0.5ml complete media. Viable cells were counted by haemocytometer using trypan blue exclusion. Serum was acquired by collecting blood from a peripheral artery using Na-EDTA coated capillaries and centrifuging at 12,000 g for 15 mins.

### Flow cytometry assessment

Lung, BAL, mLN and splenic cells were stained for flow cytometric analysis. For surface markers, cells suspensions were stained in flow cytometry buffer (PBS containing 2% fetal calf serum and 2mM EDTA). To reduce non-specific binding, cell suspensions were incubated with antibody cocktails containing anti-Fc receptor binding antibody (anti-CD16/32). Cells were extracellularly stained in antibody cocktails for 30mins at 4˚C, apart from stains containing CXCR5 which were incubated at room temperature in the dark for 1 hour. For detection of intracellular cytokines, cells were incubated with 50 ng/ml phorbol myristate acetate, 500 ng/ml inomycin and 10 μg/ml brefeldin A for 5 hours at 37 ˚C, 5% CO_2_. Cells were fixed with 1% paraformaldehyde. For detection of intranuclear transcription factors, cells were fixed and permeabilised using the Foxp3/Transcriptional factor staining buffer set (eBioscience, CA, USA) according to the manufacturer’s instructions. Cells were then washed and intracellularly stained for between 30 and 60 mins at 4˚C in permeabilization wash buffer (Biolegend, CA, USA). Flow cytometry data was acquired using an LSRII Fortessa (Becton Dickson, NJ, USA) and analysed using the FlowJo 10 software (FlowJo, OR, USA). Flow cytometry antibodies are listed in the Table S1.

### Assessment of lung function

Airway hyper-responsiveness was measured in anesthetised and tracheotomised mice in response to increasing doses of methacholine (3–100 mg/ml; Sigma-Aldrich, MO, USA) using the flexiVent system (Scireq, Montreal, Canada) as previously described^54^.

### Antibody assessment

Serum total IgE and IgG1 were measured using paired antibody ELISA sets from BD Pharmingen^TM^ (Oxford, UK). Allergen specific IgE and IgG1 levels were measured by coating plates with the appropriate allergen, adding serially diluted serum and biotinylated IgG1 or IgE (BD Pharmingen^TM^, Oxford, UK). End-point titre was calculated using baseline+2xSD based on naïve animals.

### Cytokine analysis

Lung IL-13 and IL-17A were determined using the eBioscience Ready Set Go Kit (eBioscience, CA, USA). Lung eotaxin-2 was measured using the mouse CCL24/Eotaxin-2 DuoSet ELISA (R&D systems, MN, USA). Paired antibodies for murine IL-5 (R&D Systems, Abington, UK) were used in a standardised sandwich ELISA. All ELISAs were performed according to the manufacturer’s instructions.

### Statistical analysis

Statistical significance between groups was determined using the Mann Whitney U Test and all statistical tests were performed using Prism 6 (GraphPad Software Inc, CA, USA). All *P* values ≤0.05 (*) ≤0.01 (**), ≤0.001 (***) and ≤0.0001 (****) were considered significant.

## ACKNOWLEDGEMENTS

Thanks to Professor Andrew McKenzie, University of Cambridge for gifting us the IL-13^GFP^ reporter mice. We would like to thank Catherine Simpson, Jane Srivastava and Jess Rowley from the Imperial College London, Flow Cytometry Facility. Furthermore, we want to acknowledge Lorraine Lawrence of the Research Histology Facility for carrying out the lung lobe embedding, sectioning and staining for histological analysis. This work was supported by a Sir Henry Dale Fellowship (101372/Z/13/Z) from the Wellcome Trust, UK and the Royal Society, UK to J.A.H. and a Wellcome Trust Senior Fellowship (107059/Z/15/Z) to C.M.L. In addition to a studentship from the MRC & Asthma UK Centre in Allergic Mechanism of Asthma awarded to F.I.U.

## AUTHOR CONTRIBUTION

F.I.U, J.A.H and C.M.L designed experiments. Experimental work was also carried out by F.I.U, J.A.H, C.J.P and S.A.W and analysed by F.I.U. The manuscript was written by F.I.U in collaboration with J.A.H and C.M.L who both provided feedback.

